# Active Vision in Immersive, 360° Real-World Environments

**DOI:** 10.1101/2020.03.05.976712

**Authors:** Amanda J. Haskins, Jeff Mentch, Thomas L. Botch, Caroline E. Robertson

## Abstract

Vision is an active process. Humans actively sample their sensory environment via saccades, head turns, and body movements. Yet, little is known about active visual processing in real-world environments. Here, we exploited recent advances in immersive virtual reality (VR) and in-headset eye-tracking to show that active viewing conditions impact how humans process complex, real-world scenes. Specifically, we used quantitative, model-based analyses to compare which visual features participants prioritize over others while encoding a novel environment in two experimental conditions: active and passive. In the active condition, participants used head-mounted VR displays to explore 360º scenes from a first-person perspective via self-directed motion (saccades and head turns). In the passive condition, 360º scenes were passively displayed to participants within the VR headset while they were head-restricted. Our results show that signatures of top-down attentional guidance increase in active viewing conditions: active viewers disproportionately allocate their attention to semantically relevant scene features, as compared with passive viewers. We also observed increased signatures of exploratory behavior in eye movements, such as quicker, more entropic fixations during active as compared with passive viewing conditions. These results have broad implications for studies of visual cognition, suggesting that active viewing influences every aspect of gaze behavior – from the way we move our eyes to what we choose to attend to – as we construct a sense of place in a real-world environment.

**Significance Statement:** Eye-tracking in immersive virtual reality offers an unprecedented opportunity to study human gaze behavior under naturalistic viewing conditions without sacrificing experimental control. Here, we advanced this new technique to show how humans deploy attention as they encode a diverse set of 360º, real-world scenes, actively explored from a first-person perspective using head turns and saccades. Our results build on classic studies in psychology, showing that active, as compared with passive, viewing conditions fundamentally alter perceptual processing. Specifically, active viewing conditions increase information-seeking behavior in humans, producing faster, more entropic fixations, which are disproportionately deployed to scene areas that are rich in semantic meaning. In addition, our results offer key benchmark measurements of gaze behavior in 360°, naturalistic environments.

## Introduction

Constructing a sense of place in a complex environment is an active process. Humans actively sample their sensory environment to understand their surroundings and gain information relevant to their behavioral goals (Hayhoe, 2017; Hayhoe and Matthis, 2018). Yet, much of what we know about how people encode real-world environments comes from computer-based paradigms that severely limit participants’ active affordances. In this context, the participant’s behavioral repertoire is limited to eye movements, and the displayed environment is typically limited to a single field of view. In contrast, everyday visual environments are actively explored. We gain rich information about a place by shifting our eyes, turning our heads, and moving our bodies. This is because real-world scenes are immersive, extending 360º around us and beyond any single field of view. How does scene understanding unfold in immersive, active viewing conditions?

It has long been understood that active viewing conditions impact perceptual processing. Neurons in early stages of the visual system are sensitive to the distinction between self- and world-generated motion under conditions that are carefully matched for retinal stimulation and attentional engagement (Goldberg and Wurtz, 1972; Troncoso et al., 2015). Further, active vision is thought to be necessary for typical visual development: even basic functions, such as depth perception and contrast sensitivity, suffer when animals are denied self-motion, but are passively exposed to equivalent visual environments (Held and Hein, 1963). Studies in humans also suggest that perceptual systems differentially represent stimuli that are encountered via active vs. passive viewing (Hayhoe, 2017). For example, an object’s spatial location is better recalled when it has been actively reached for rather than passively moved toward (Trewartha et al., 2015). Yet, to date, no studies have explored how active viewing conditions impact the processing of complex visual stimuli, such as real-world scenes.

Here, we used a novel experimental design to study real-world scene processing during active and passive viewing conditions. We exploited recent developments in virtual reality (VR) to immerse participants in real-world, 360º scenes. Meanwhile, we monitored participants’ gaze using in-headset eye-tracking as they explored these environments, revealing which regions they prioritized over others. In one condition, participants explored scenes from an active, first-person perspective. In the other condition, scenes were passively displayed to participants while they were head-restricted in a chin rest. In both conditions, diverse, real-world scenes were displayed with the same wide-angle field of view (100 DVA), and participants were exposed to comparable portions of the display over the course of the trial. Thus, this paradigm enabled us to perform quantitative, in-depth comparisons of gaze behavior and attentional deployment as subjects encoded novel, real-world scenes during active vs. passive exploration.

Our central hypothesis was that active viewing conditions would increase viewers’ exploratory, information-seeking behavior in a real-world scene. We tested this by measuring the degree to which participants’ overt attention was dominantly predicted by the spatial distribution of scene features that are semantically informative (e.g., objects, faces, doors) (Henderson and Hayes, 2017; Henderson et al., 2018), vs. scene regions that are rich in salient visual features (e.g., luminance, contrast, color, and orientation) (Itti and Koch, 2000; Greene and Oliva, 2009). Previous studies have shown that these information sources compete for participants’ top-down vs. bottom-up attention, although attention is predominantly predicted by the distribution of semantic information (Henderson and Hayes, 2017, 2018).

In brief, we observed that participants’ attention was dominantly guided by semantic meaning as compared low-level visual features in both active and passive conditions, replicating previous findings (Henderson and Hayes, 2017, 2018). Crucially, this effect tripled during active viewing, reflecting an increase in signatures of top-down attentional guidance when participants were free to actively view their environment. Further, in service of this information-seeking behavior, active viewers made shorter, more exploratory fixations than passive viewers. These results show that active viewing influences every aspect of gaze behavior, from the way we move our eyes to what we choose to attend to.

## Methods

### Participants

Eighteen adults participated in the main experiment (thirteen females; mean age 22 years +/−3.73 STD). Three additional participants completed a pilot study designed to test eye-tracker accuracy and precision (Supplemental Figure 1). Participants were recruited based on 1) having normal or contact-corrected vision and no colorblindness, 2) having no neurological or psychiatric conditions, and 3) having no history of epilepsy. Written consent was obtained from all participants in accordance with a protocol approved by the Dartmouth College and Massachusetts Institute of Technology Institutional Review Boards.

**Figure 1.**
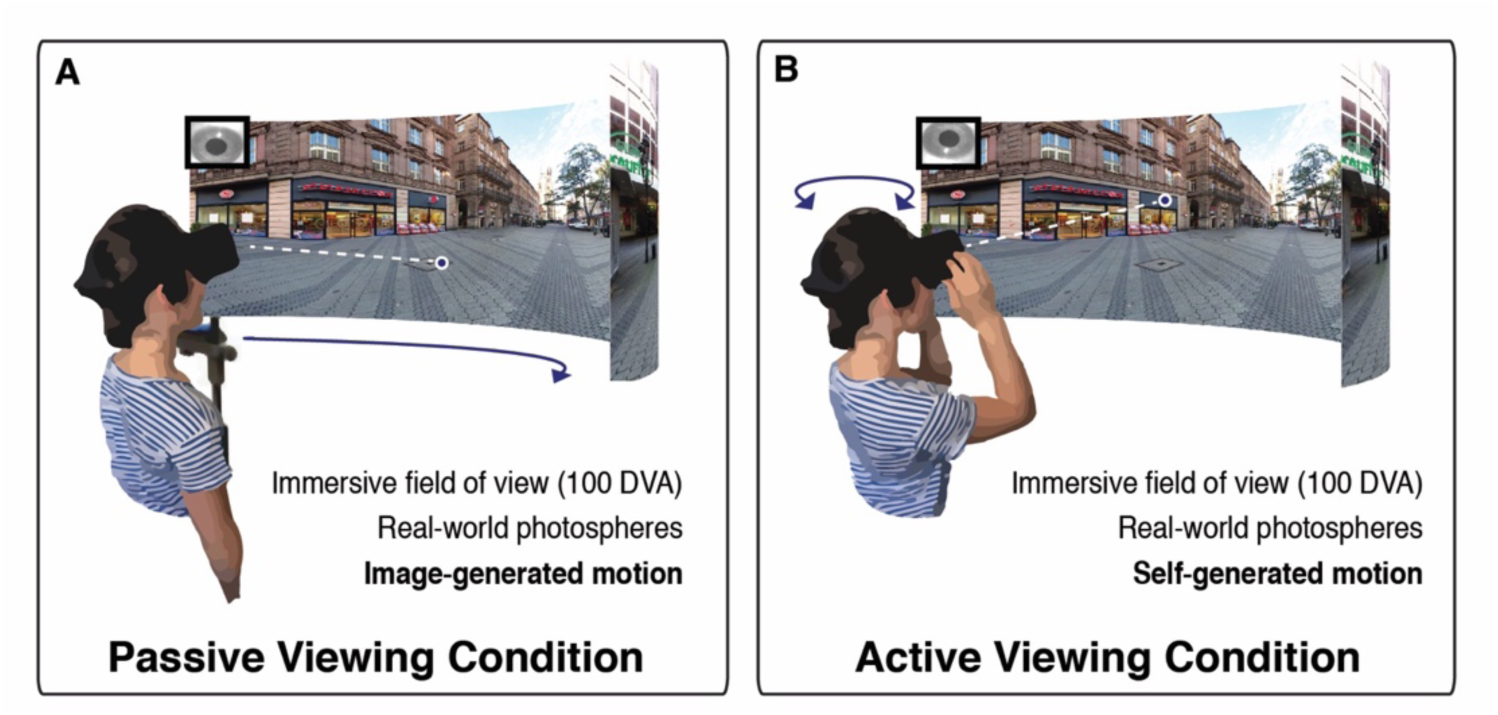
Passive viewing condition vs. active viewing condition. On each trial, participants viewed immersive, 360° real-world scenes via a headmounted VR display while gaze position was monitored using an in-headset eye-tracker**. A)** In the passive condition, participants were head-restricted using a chin rest, and scenes panned across the display. **B)** In the active condition, participants explored scenes from a first-person perspective through movement of their eyes, head, and body.

### Stimulus and headmounted display

Stimuli consisted of 360° “photospheres” of real-world scenes, sourced from an online photo sharing website (www.flickr.com). Photospheres depicted a diverse set of indoor and outdoor settings with content including people and objects. Each photosphere was applied to a virtual environment built in Unity version 2017.3.1f1 (www.unity3d.com) and integrated with a headmounted display (Oculus Rift, Development Kit 2, www.oculus.com, low persistence OLED screen, 960 × 1080 resolution per eye; ~100 degree field of view; 75 Hz refresh rate).

### Eye-tracker specifications and accuracy

A monocular, in-headset eye-tracker (Pupil Labs: 120 Hz sampling frequency, 5.7 ms camera latency, 3.0 ms processing latency; 0.6 visual degrees accuracy, 0.08 visual degrees precision) continuously monitored the position of participants’ right eye during scene viewing. Eye movements were recorded using custom scripts written in C# for Unity. Refer to Supplemental Figure 1 for observed accuracy and precision of Pupil Labs eye-tracker at varying eccentricities.

### Experimental Design

On each trial of the experiment (40 trials), participants were presented with a photosphere via the headmounted display (HMD). Participants were instructed to “fully and naturally explore each scene”. Participants were given a break after every 10 scenes, after which the eye-tracker was recalibrated.

There were two viewing conditions in this experiment: active and passive (Figure 1). In both conditions, the stimulus was presented via the HMD and each trial lasted for 20s. During the active condition, participants stood while wearing the HMD and actively explored the photosphere via self-directed eye movements and head turns. In contrast, during the passive condition, participants’ heads were fixed in a chin rest and the scene panned across the screen, rotating 360° at a constant velocity (22°/ second). In order to prevent the sensation of motion sickness, rotational velocity gradually ramped up and down during the first and last two seconds of each passive trial.

There were 40 total stimuli included in the experiment. Each participant viewed 20 stimuli in each of the two conditions with condition assignment randomized for each trial and participant. Conditions were blocked, but condition order was counterbalanced across participants. The initial rotation angle of each scene was held constant across participants.

### Practice Trials and Calibration Routine

There were three phases to the experiment: practice, calibration, and experimental trials. During the practice phase, participants performed two active condition trials. This ensured that participants had acclimated to the virtual environments prior to starting the experiment. Following the practice phase, participants performed a 14-point calibration routine in order to validate eye-tracking accuracy. Participants repeated the calibration routine after every 10 experimental trials.

After each trial in the Experimental Phase, participants returned to a virtually rendered home screen where they were instructed to take a break. Upon advancing from the home screen, participants were presented with a pre-trial fixation screen with a target at screen center. Participants were instructed to fixate on the target so that gaze drift could be assessed. If significant drift (> 5 degrees visual angle) was detected, a recalibration routine was performed.

### Eye-tracking data analysis

Raw *x (*and *y)* gaze points were converted from normalized screen coordinates to DVA using the following equation(Ehinger et al., 2019):

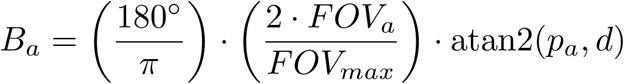

where, *B*_*a*_ denotes the azimuth (or elevation) angle of gaze position in visual degrees relative to screen center, *FOV*_*a*_ denotes the field of view of the x (or y) dimension of the HMD as a proportion of the largest dimension of the HMD (i.e., 100 DVA), *p*_*a*_ denotes the gaze position in normalized screen coordinates, and *d* denotes the distance in Unity units that places the participant at the origin of the spherical eye-tracking coordinate system. Next, gaze coordinates were rectified with head position (pitch, yaw, roll), and transformed into latitude and longitude positions on a sphere (spherical degrees):

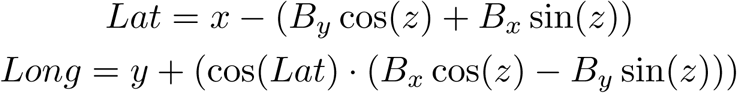

Here, *x* denotes pitch in spherical degrees (with up being negative and down being positive), *y* denotes yaw in spherical degrees, and *z* denotes roll in radians. *B*_*x*_ and *B*_*y*_ denote gaze point distance from screen center in visual degrees.

Within each trial, a gaze point was labelled as invalid if: 1) it fell outside the field of view (i.e., greater than 50° from screen center in either the *x* and/or *y* direction), 2) pupil detection confidence was low (i.e., below 50 percent), or 3) no data was collected (e.g., during a blink). Trials with more than 75 percent of points labelled as invalid were excluded from the analysis.

### Defining fixations

Next, to determine fixations, the orthodromic distance and velocity was calculated between consecutive gaze points. Specifically, the mean absolute deviation (MAD) (Voloh et al., 2019) in gaze position was calculated within a seven-sample sliding window (~80ms) and potential fixations were defined as windows with a MAD less than 50°/s (Peterson et al., 2016). Potential fixations were concatenated if two group centroids were displaced by less than 1° and the two potential fixations occurred within 150ms of each other. Fixations with durations shorter than 100ms were excluded (Wass et al., 2013; Peterson et al., 2016). To standardize fixation density maps across conditions, fixations in the active condition were excluded if they fell beyond the region displayed during the passive condition (27.9 percent of the spherical scene; Figure 2D). Given the infrequency with which participants looked at the upper and lower poles of a scene, only 12.2 percent +/−1.29 STE of fixations were discarded from the active condition. Additionally, because the rotational velocity was changing at the beginning and end of each passive condition trial, fixations made during the first and last two seconds were excluded from both active and passive conditions.

**Figure 2.**
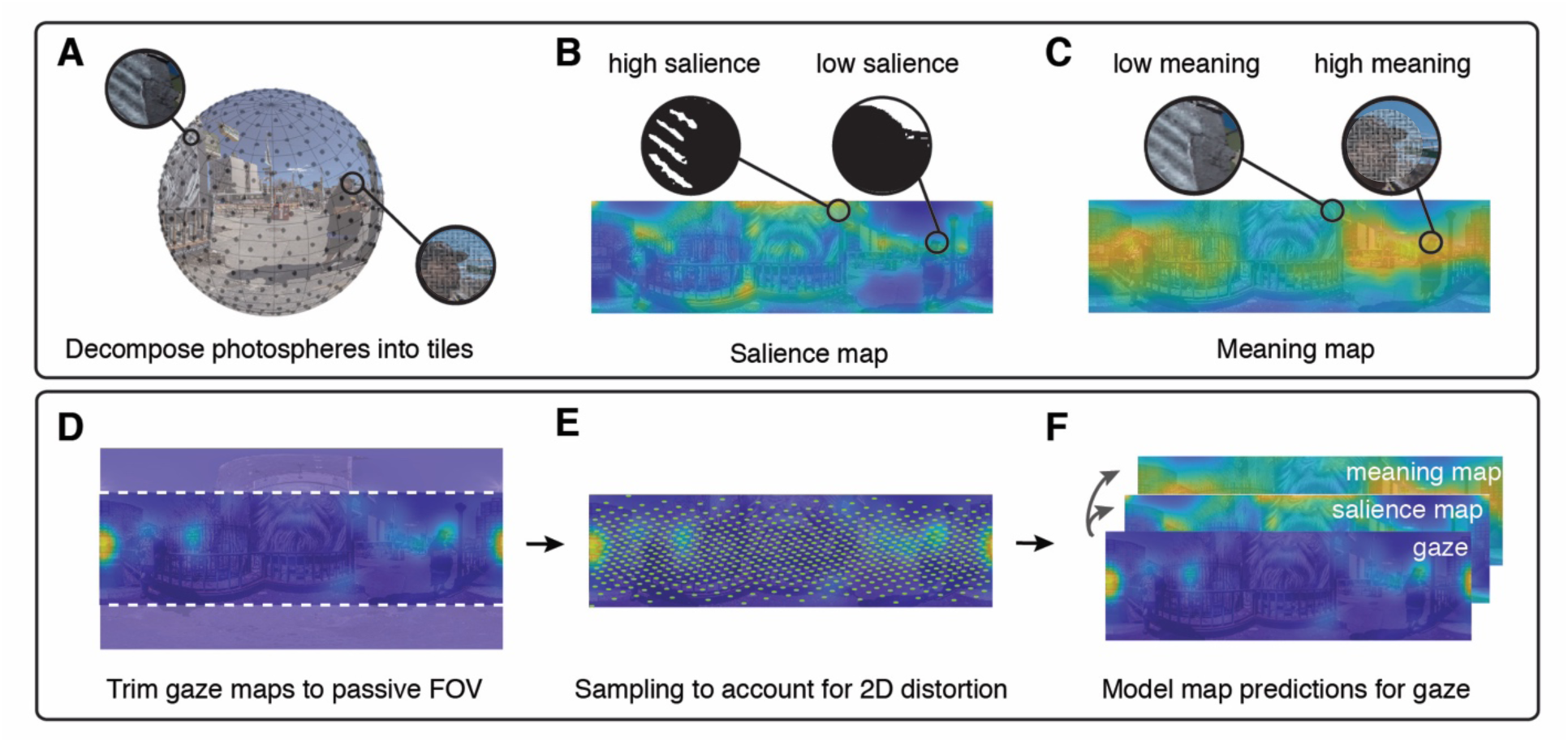
Comparing salience and meaning maps to gaze behavior. **A)** Each photosphere was first decomposed into smaller undistorted image tiles. Next, we created two models of the content in each real-world environment. **B)** “Salience maps” were generated by modeling low-level visual features for each tile using the GBVS Toolbox(Harel et al., 2007). Each tile was then projected onto a two-dimensional salience map. **C)** “Meaning maps” were generated via online participants who rated the semantic content, or “meaning” of each image tile. Each tile’s rating was then projected onto a two-dimensional meaning map. **D)** Group gaze maps were trimmed (vertically) to match the passive condition field of view. **E)** Points are sampled evenly on a sphere and used to account for photosphere distortion in two-dimensional maps. **F)** A linear mixed effects model was used to compare the degree to which each model predicted attentional guidance in our two conditions.

### Fixation density map generation

We next characterized the spatial distribution of fixations on each trial. To generate two-dimensional fixation density maps, we first plotted the fixations for all subjects in a given scene and condition in equirectangular space. The resulting fixation maps were smoothed with a variable-width gaussian filter (“modified gaussian” (John et al., 2019)) to account for distortions of the equirectangular image at shorter latitudes (i.e., approaching the poles). Specifically, the width of the gaussian filter is scaled by the latitude of the gaze point using the following equation:

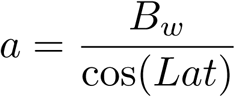

where the width of the filter, *a*, at a given latitude (*Lat*) has been scaled from the base filter width (*B*_*w*_) applied at the equator.

### Gaze map generation

Finally, we generated duration-weighted fixation density maps. In order to prevent extreme fixation durations made by individual subjects from exerting an outsized impact on the group gaze maps, fixations with durations above the 95^th^ percentile were reduced to the 95^th^ percentile value (Henderson and Hayes, 2017, 2018). Additionally, each individual subject’s fixations were normalized on a scale from 0.1 to 1.

### Quantifying central tendency

A routine observation in fixed display studies is the tendency for fixations to be disproportionately allocated at the center of a scene (Bindemann, 2010). “Central tendency” has been employed as a metric of visual exploration, where fixations that are less centrally tending are considered more exploratory (Gameiro et al., 2017). Given the tendency for viewers to fixate near the equator in VR (Sitzmann et al., 2018), “equator bias” was calculated per condition by averaging the distance of each fixation from the equator in the *y* dimension only.

### Quantifying entropy

The entropy of the resulting fixation density map was calculated using the following equation (Açik et al., 2010):

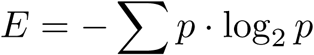

where p contains the fixation density map’s histogram counts. Because entropy estimates can be impacted by small sample sizes (Wilming et al., 2011), and because an uneven number of fixations were made across conditions for any single scene, we applied a bootstrapping technique to estimate entropy. Across 100 iterations, we randomly sampled 24 fixations from each participant within each scene in a given condition. The target of 24 fixations was chosen in proportion to previous studies analyzing the entropy of nine fixations per 6s trial (Kaspar et al., 2013; Gameiro et al., 2017).

### Salience map and meaning map generation

Our central hypothesis was that active viewing conditions would increase exploratory, information-seeking behavior in a real-world scene. To test this hypothesis, we constructed two models of the visual content in each environment. First, we computed a traditional "salience map”, which reflects the distribution of low-level visual features in a scene (e.g., contrast, color, orientation, etc.) (Harel et al., 2007). Prominent low-level visual features in a scene are known to predict a significant portion of gaze behavior (Itti and Koch, 2000; Parkhurst et al., 2002). However, as attention is drawn on the basis of visual salience, rather than semantically meaningful scene regions, the degree to which such maps predict gaze behavior is not related to information-seeking behavior, thus providing a good baseline model for our hypothesis. Second, we computed a “meaning map” for each scene, which reflects the distribution of high-level semantic features in an environment (e.g., faces, objects, doors, etc.) (Henderson and Hayes, 2017, 2018). “Meaning maps”, a recently proposed “conceptual analogue” of salience maps (Henderson and Hayes, 2017, 2018), reflect the spatial distribution of features that are relevant to understanding the semantic content and affordances available to the viewer in a scene. Recent studies have shown that attentional deployment in novel scene images is dominantly predicted by the spatial distribution of scene features that are semantically informative (meaning maps) as compared with low-level features (salience maps) (Henderson and Hayes, 2017, 2018).

Salience maps were generated using the Graph-Based Visual Saliency (GBVS) Toolbox (Harel et al., 2007). Each photosphere was uniformly sampled and decomposed into a set of 500 square tiles, each with a diameter of 7.5° (Figure 2A). The GBVS model with default feature channels (i.e., color, intensity, orientation) was applied to each tile, which was then projected back to its position in the equirectangular image (Figure 2B). Salience maps were smoothed using the variable-width gaussian filter applied to gaze maps.

To generate meaning maps, we applied the procedures described by Henderson and Hayes (Henderson and Hayes, 2017, 2018) to 360° scenes. Each photosphere was uniformly sampled at both coarse (100 points) or fine (500 points) spatial scales and decomposed into sets of partially overlapping circular tiles with diameters of 20.6 spherical degrees or 7.5 spherical degrees, respectively. Scene tiles were produced by generating a rectilinear projection (1100×1100 pixels) around each point sampled on the sphere. Each coarse tile was down-sampled to match the resolution of fine tiles, resulting in a diameter of 150 pixels for all scene tiles. The full scene tile stimulus set contained 20,000 unique fine-scale tiles and 4,000 unique coarse-scale tiles, for a total of 24,000 scene tiles.

A total of 1,879 participants on Amazon Mechanical Turk rated scene tiles on a 6-point Likert scale (very low, low, somewhat low, somewhat high, high, very high). Participants were instructed to rate the content of each scene tile based on how “informative or recognizable” it was. Participants were first given examples of two low-meaning and two high-meaning tiles, followed by a set of four practice trials to ensure understanding of the task. The practice trials contained two examples expected to score on the low-meaning side of the scale (1-3) and two examples expected to score on the high-meaning side of the scale (4-6). Seventy-nine participants were excluded based on the results of these diagnostic practice trials.

Each participant rated 40 tiles (20 of each spatial scale). Participants rated exactly one tile from each of the 40 scenes, and participants were prevented from completing the online experiment more than once. In total, the experiment took approximately 3 minutes and participants were compensated for completing the study.

Each tile was rated by three participants, and responses were averaged to produce a “meaning” rating for each tile (Figure 2C). The average rating for each tile was then plotted at its center coordinate and smoothed using the variable-width gaussian filter. This process was completed at both spatial scales, and the average of these two maps was used as the scene’s final “meaning map”.

### Spherical sampling of equirectangular maps

To account for the distortion imposed by equirectangular map projections, which disproportionately represent scene regions at the poles, we sampled (N = 100,000) points uniformly on a sphere, projected those indices onto each equirectangular map (i.e., gaze, salience, meaning, and equator), and used the sampled values at these indices for analyses of spatial attention (Figure 2E) (Gutiérrez et al., 2018). As a result, each individual index, or location, in an equirectangular map was treated as a separate observation (n = 6,164,000) in statistical analyses.

### Statistical analyses

To compute the relative contributions of visual salience and semantics in predicting gaze behavior, we built a linear mixed effects model using the lme4 package in R (Bates et al., 2015). We included viewing condition (i.e., active vs. passive) and feature maps of scene content (i.e., salience, meaning, and equator map) as fixed effects and individual scenes as random effects. Specifically, for each scene, the model predicted the degree to which each feature map predicted the gaze map value at each location (each index from the N=100,000 points uniformly sampled around the spherical image). Because gaze maps were generated at the group level (Henderson and Hayes, 2017, 2018), individual subjects were not included as random effects in the model. Two-way interactions (i.e., salience by condition, meaning by condition) and the three-way interaction between salience, meaning, and condition were analyzed.

## Results

To test whether active viewing conditions modulate attentional guidance in real-world scenes, we directly compared eye movements while participants viewed immersive, 360° environments in the two experimental conditions. We observed high-quality eye-tracking in the HMD, comparable with that reported in fixed display studies at screen center (accuracy: 0.79 DVA, precision, 0.11 DVA; see Supplemental Figure 1).

We specifically hypothesized that active viewing conditions would increase exploratory, information-seeking behavior, and therefore increase the degree to which meaning-maps predict gaze behavior. One of the few existing observations from eye-tracking in VR is the tendency for viewers to fixate near the equator of 360° scenes (Sitzmann et al., 2018) (a 3D equivalent of “center bias”). Therefore, to test our hypothesis, we first generated a map of the equator to serve as a baseline prediction for 360º viewing behavior. Then, for each condition, we compared the additional predictive contribution of each map of environmental content (salience map, meaning map) using a linear mixed-effects model. Specifically, we included three spatial maps (i.e., salience maps, meaning maps, and our baseline map of the equator) and viewing condition as fixed effects and individual scenes as random effects in the model. All results are summarized in Table 1.

**Table 1:** Linear mixed effects model results summary.

|  | **numDF** | **denDF** | **F value** | **p value** |
| --- | --- | --- | --- | --- |
| condition | 1 | 6163960 | 8818.25 | < 0.001 |
| meaning | 1 | 6163907 | 71586.82 | < 0.001 |
| saliency | 1 | 6163982 | 19920.19 | < 0.001 |
| condition:meaning | 1 | 6163960 | 21579.7 | < 0.001 |
| condition:saliency | 1 | 6163960 | 212.22 | < 0.001 |
| meaning:saliency | 1 | 6163997 | 64200.96 | < 0.001 |
| condition:equator | 2 | 6163980 | 485605.49 | < 0.001 |
| condition:meaning:saliency | 1 | 6163960 | 71.36 | < 0.001 |

Overall, we observed that overt attention in real-world scenes primarily reflects information-seeking behavior in both active and passive conditions, confirming previous results (Henderson and Hayes, 2017, 2018). Both salience and meaning maps significantly predicted participants’ overt attention (salience estimated marginal effect: 0.15 +/− 0.03 STE, CI: [0.09,0.21]; *p* < 0.001; meaning estimated marginal effect: 0.31 +/−0.03 STE, CI: [0.25, 0.36]; *p* < 0.001). However, in both active and passive conditions, meaning was significantly more predictive of which scene regions participants explored than salience (meaning:salience interaction: *p* < 0.001; post-hoc corrected t-tests for meaning vs. salience: *p* < 0.001 for both conditions).

Critically, however, this advantage for meaningful scene regions nearly tripled in the active as compared with the passive condition (salience*meaning*condition: *p* < 0.001; Figure 3). Post-hoc analyses revealed that the estimated marginal effect of meaning (i.e., its predictive contribution, holding other factors constant) was significantly greater for active as compared with passive viewers (passive: 0.25 +/−0.03 STE; active: 0.31 +/−0.03 STE; *p* < 0.001). Conversely, the estimated marginal effect of salience was greater in the passive condition than in the active condition (passive estimate: 0.20 +/−0.03 STE; active estimate: 0.15 +/−0.03 STE; *p* < 0.001). Taken together, these results show that active viewing conditions specifically increase attentional allocation to semantically relevant regions of a visual environment.

**Figure 3.**
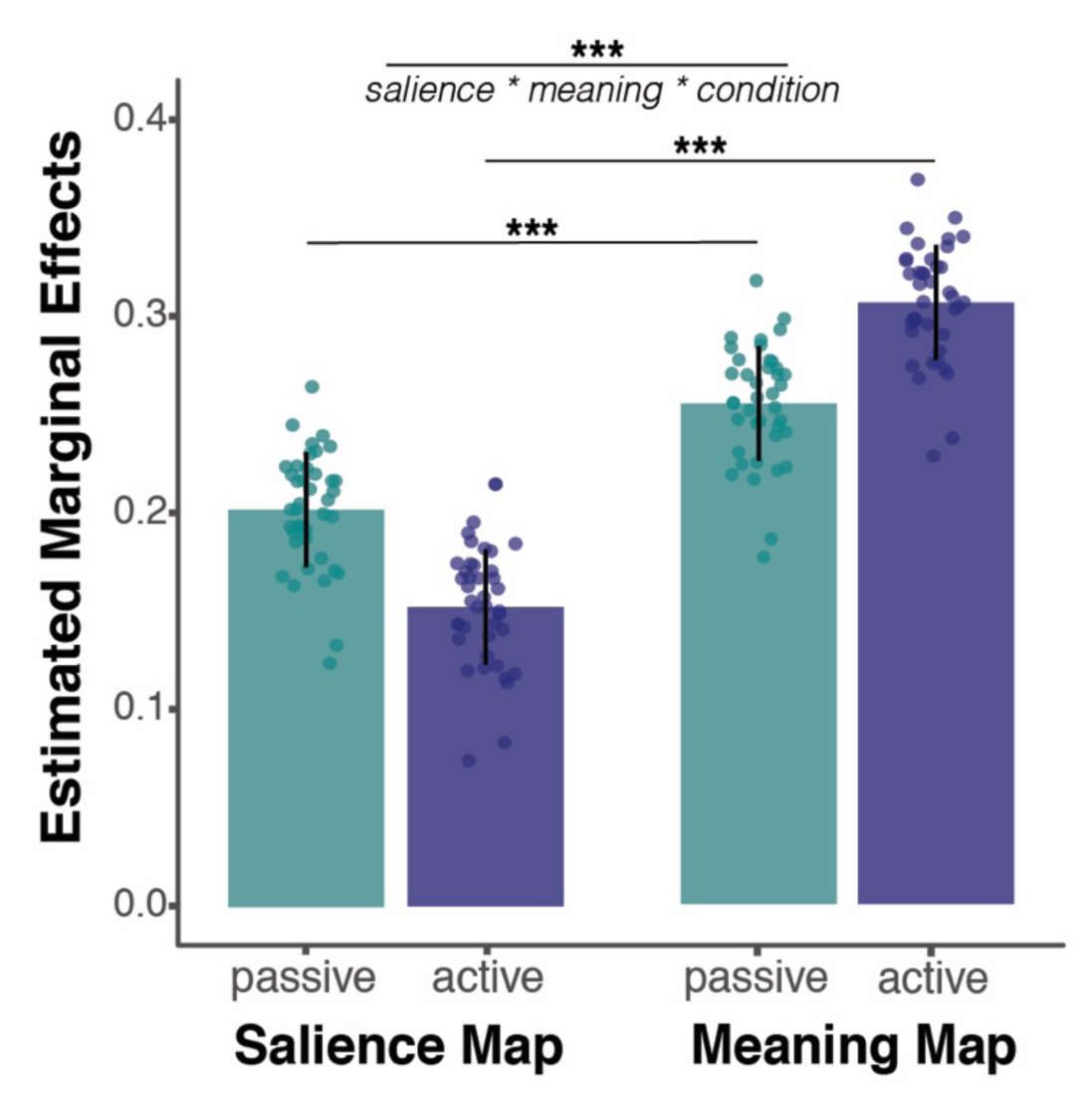
Active viewing increases top-down attentional allocation. Estimated marginal effects of salience and meaning maps on predicting overt attention in each condition (active vs. passive). We found that viewing condition (active vs. passive) significantly modulated gaze behavior (salience*meaning*condition: *p* < 0.001). Specifically, active viewers disproportionately directed their attention to meaningful, over salient, scene regions. Individual points represent random item effects (i.e., individual scenes). Error bars represent prediction intervals (+/− 1 STE). *** denotes *p*< 0.001.

Control analyses confirmed that these results could be attributed to active affordances rather than passive viewers’ limitations. First, we found that the disproportionate advantage for meaning-guided attention in the active condition remained significant even after restricting our analysis to the fields of view containing regions ranked in the top 50th percentile for meaning, thereby eliminating less meaningful scene regions that passive viewers were required to scan as a result of the panning image presentation (Supplemental Table 1). Second, we found that our results held even when accounting for neighboring fixations in the active condition whose combined duration exceeded the time any given scene region would have been displayed to a participant in the passive condition (5s). In fact, we found that participants in the active condition rarely fixated within a region for longer than it would have been displayed during the passive condition (< 3% of fixations), even when accounting for return fixations. Thus, as predicted, in the active condition, when participants were free to move their head and body, participants’ attention was disproportionately directed towards semantically relevant regions of a visual environment.

We further characterized gaze behavior during active viewing using two measures of the spatial distribution of attention that are independent of individual scene content: 1) deviation from center bias and 2) entropy. Studies of visual attention using traditional fixed displays commonly observe a bias to fixate near the center of an image (Tatler, 2007; Bindemann, 2010). Although the precise source of this bias is disputed (Parkhurst et al., 2002; Schumann et al., 2008; Tseng et al., 2009), deviation from “center bias” has been used to describe the degree of visual exploration (Gameiro et al., 2017). On average, we found that fixations made in the active condition had less center bias, or were further from the equator (Sitzmann et al., 2018), (17.13 degrees +/−0.54 STE) than fixations made in the passive condition (12.53 degrees +/−0.23 STE) (*t*(39) = 9.98, *p* < 0.001; Figure 4). Of course, this result could reflect a systematic, non-central bias (e.g., active viewers could have routinely looked toward the poles), rather than more exploratory gaze behavior *per se*. To address that possibility, we used a second measure, entropy, a measure of homogeneity (or lack thereof) in the probability distribution of fixations (Açik et al., 2010), to test whether any systematic biases occurred in each viewing condition. We found that gaze behavior in the active condition was more entropic (3.49 +/−0.07 STE) than gaze in the passive condition (2.88 +/−0.02 STE) (*t*(39) = 8.63, *p* < 0.001). Taken together, these results further demonstrate that active viewers prioritize rapid exploration of new scene regions that are rich in semantic content, relative to passive viewers.

**Figure 4.**
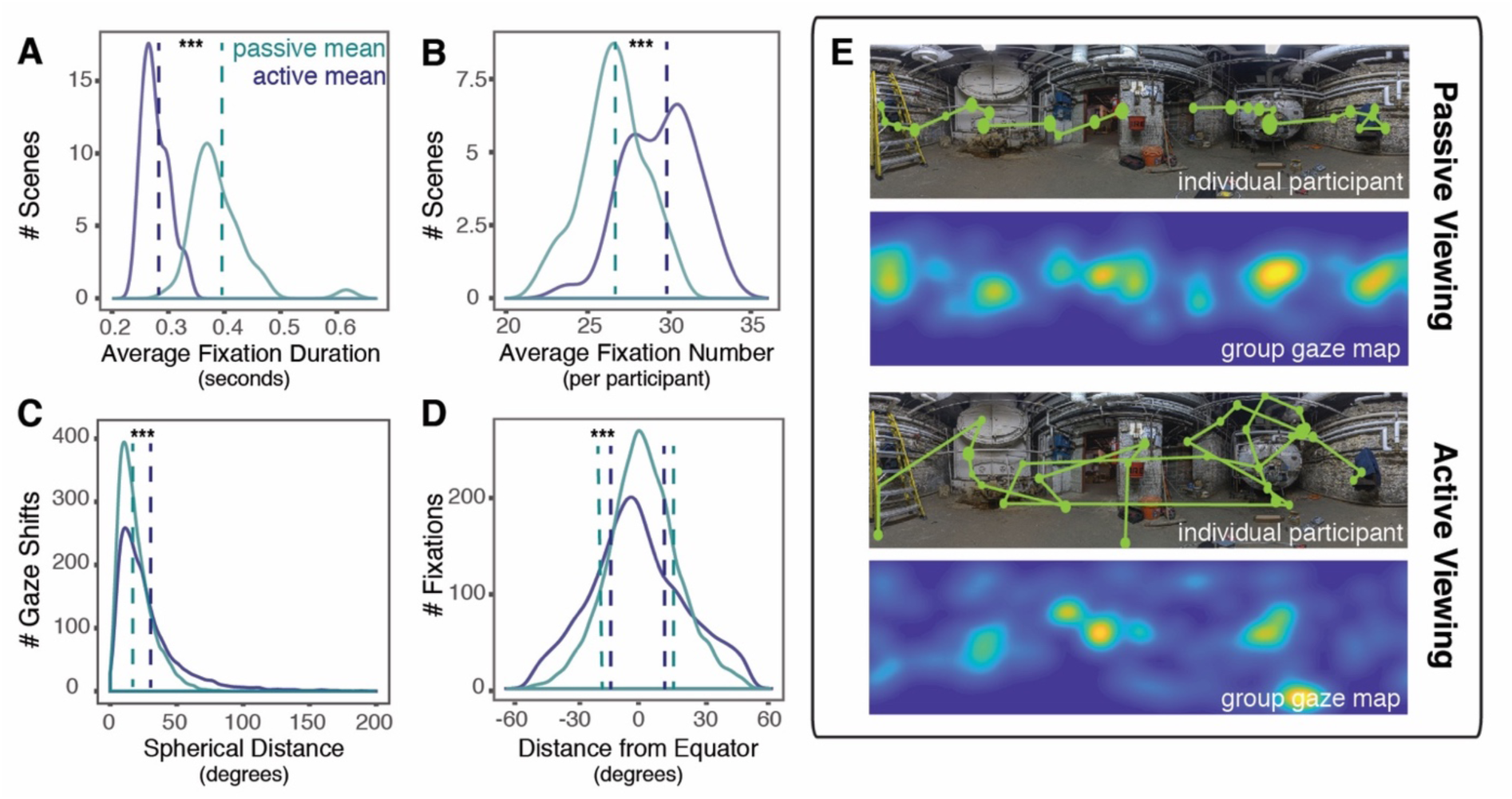
Active viewing impacts eye movements. A) Relative to fixations made in the passive viewing condition, fixations in the active viewing condition were shorter B) and more frequent. C) Gaze shifts in the active viewing condition were also larger, and D) the spatial distribution of gaze in the active viewing condition was less centrally tending. E) Sample duration-weighted fixations and gaze shifts made by a single participant (top) and group fixation map for participants (bottom) per condition. *** denotes *p* < 0.001.

Finally, we sought to characterize the eye movements participants made in service of this increase in information-seeking behavior (Figure 4). Participants made shorter (0.27s +/−0.004 STE) and more frequent (29.45 +/− 0.34 STE) fixations in the active as compared with the passive condition (size: 0.39s +/− 0.008 STE, *t*(39) = − 14.52, *p* < 0.001; frequency: 26.60 +/− 0.31 STE, *t*(39) = 7.53, *p* < 0.001. Further, active viewing conditions impacted the magnitude of gaze shifts (i.e., the combined movement of eyes and head between two fixations) participants made while exploring their environment. Gaze shifts were significantly larger in the active condition (29.17 DVA +/− 0.48 STE) as compared with the passive condition (19.08 DVA +/− 0.23 STE: *t*(39) = 23.11, *p* < 0.001). Indeed, it was not uncommon for active viewers to make gaze shifts as large as 150 DVA, exceeding previous saccade length estimates from fixed display studies (5-10 DVA) by an order of magnitude (Land and Hayhoe, 2001; Tatler et al., 2006). Notably, our headmounted display subtended a wider field of view (100 DVA) than typically afforded by fixed display studies. Importantly, this wide display was matched for both active and passive conditions; thus, the differences we observed cannot be attributed to the size or content of our stimuli, which is known to impact the size of gaze shifts (Von Wartburg et al., 2007). Instead, our results demonstrate that active viewing conditions fundamentally impact the size of gaze shifts a viewer will routinely choose to make when exploring a real-world visual environment: people make faster fixations and larger gaze shifts, once more suggesting more exploratory gaze behavior.

## Discussion

Our results provide novel insights into the process by which we actively construct representations of immersive, real-world visual environments. We found that, when participants are unconstrainted and free to choose their field of view, their behavior is most guided by meaningful, semantic properties of the environment, as compared with passive viewers. Moreover, in service of this information-seeking behavior, active viewers employ shorter, more entropic fixations. All in all, we demonstrate that active viewing conditions impact nearly all features of gaze behavior, from gaze mechanics (how we move our eyes) to gaze dynamics (what we choose to attend to). These findings lay the foundation for future studies of gaze behavior in immersive, real-world environments using VR.

The recent development of eye-tracking in immersive VR is a critical advance for investigating real-world scene processing in the context of natural behavioral affordances. VR frees the subject from the limitations typically imposed by head-restricted display studies, allowing for eye, head, and body movements (Hayhoe, 2017). Moreover, it does so without sacrificing experimental control, allowing for the presentation of a diverse set of stimuli within precise trial structures. Such stimulus diversity and controlled presentation is an essential ingredient for quantitative, model-based insights into active human gaze behavior. This opportunity for experimental control and diverse stimulus presentation stands in contrast to mobile eye-tracking paradigms, where participants might only traverse a single, extended environment (e.g., a university campus) in which the low-level visual features (e.g., the lighting/contrast) as well as high-level visual features (e.g., people walking down the street) inevitably vary between participants. Previous approaches comparing active versus passive viewing have relied on a combination of fixed-display and mobile paradigms (Foulsham et al., 2011; Peterson et al., 2016), which inherently differ in terms of task demands (e.g., watching a video vs. navigating) and displayed field of view. All in all, eye-tracking in immersive VR is an exciting opportunity to gain insight into active visual cognition.

Our findings have multiple implications for active, real-world vision. Recent studies have proposed a dominant role for semantically meaningful scene regions in guiding attention even during passive viewing of fixed-display images, suggesting that gaze behavior primarily reflects high-level information-seeking priority as scene understanding unfolds (Henderson and Hayes, 2017, 2018; Henderson et al., 2018). Our results extend these findings in three key ways. First, we show that semantically relevant features guide attention in immersive, naturalistic environments. This is an important demonstration if semantically guided attention is indeed a feature of real-world vision. Second, we show that active viewing conditions increase the advantage for semantics over low-level visual salience in guiding gaze behavior. Again, this finding has important implications for real-world vision; our results suggest that when participants are free to seek out information and choose their field of view, their behavior is most guided by meaningful, semantic properties of the scene. Finally, our findings provide key benchmark measurements of gaze behavior in diverse, real-world scenes during active viewing conditions, demonstrating that active viewers make quick, entropic fixations and shift their gaze nearly twice per second.

We attribute the observed increase in exploratory, information-seeking behavior in active viewing conditions to differences in the *affordances* available to active viewers, not differences in their *action goals*. Our experiment manipulated self- vs. image-generated motion, but not participants’ task, a factor long understood to impact gaze behavior (Yarbus, 1967; Ballard and Hayhoe, 2009; Kollmorgen et al., 2010). Participants in the active condition had no objective to physically interact with meaningful scene regions, such as objects; nor was this a possibility. Yet, access to a naturalistic behavioral repertoire – a broader capacity for self-generated action – nonetheless impacted how participants moved their eyes and deployed their attention. Our results are consistent with theoretical models linking perceptual processing and motor plans in a perception-action cycle (Wolpert and Landy, 2012), in which perceptual processes depend on movement states (Chiappe et al., 2010; Maimon et al., 2010; Jung et al., 2011; Matthis et al., 2018) and the role of vision is to provide evidence to satisfy behavioral goals (Hayhoe, 2017). These findings have important implications for future work investigating cognitive processes where motor goals are often hampered by less naturalistic paradigms, such as scene perception, spatial memory, and even social inference.

Of course, our experimental approach also has drawbacks. The design of our passive condition was limited by the known tendency for image-generated motion to induce participant motion sickness (Pan and Hamilton, 2018). As a result, we opted to implement a passive condition that slowly revealed the panoramic environments to participants, giving passive viewers the opportunity to explore the same fields of view as active viewers, but did not provide an exact match for the sequence of moment-to-moment fields of view that an active viewer would have taken in a scene. We do not think that these limitations significantly impacted our results. Both semantic and salient scene features were equally distributed around our panoramic scenes, participants in both conditions were given an equal amount of time to explore each environment, and our control analyses demonstrate that active viewers rarely, if ever, dwelled on any portion of the panoramic scene for longer than it would have been displayed to a passive viewer. Thus, the disproportionate advantage for meaning over salience in predicting gaze behavior during active viewing conditions can be attributed to which scene features a participant chose to attend to in any field of view, rather than which fields of view active vs. passive viewers sampled during a trial. In our active condition, we were limited by the challenge of self-embodiment (Pan and Hamilton, 2018): participants could move their eyes, head, and body, but their movements were not paired with the typical visual experience of seeing one’s body. It is possible that our results would strengthen if the perception of self-embodiment were afforded to active viewers, as in real-world viewing conditions. Finally, given the free viewing nature of our experiment, we cannot rule out the possibility that our effects are mediated by differences in attentional engagement between our two conditions, particularly given that VR was a novel experience for most participants.

In sum, our results provide a window into active vision during first-person exploration of immersive, real-world scenes. These findings bring into focus the importance of naturalistic behavioral affordances for studies of visual cognition, and also raise important question for future research. For example, how do differences in attentional deployment during active vision impact subsequent memory of immersive environments, or the body-based representation of such environments (Robertson et al., 2016; Huffman and Ekstrom, 2019)? How might attentional markers of clinical conditions, such as autism (Robertson et al., 2013; Wang et al., 2015), be altered by natural behavioral affordances? All in all, we show that active viewing influences every aspect of gaze behavior, from the way we move our eyes to what we choose to attend to.

## Acknowledgements

This study was supported by a grant from the Nancy Lurie Marks Family Foundation to C.E.R and the donation of a Titan V GPU by the NVIDIA Corporation to C.E.R. Thanks to Michael Cohen for insightful comments on the manuscript. Thanks also to George Wolford, Anna Mynick, Nancy Kanwisher, Brad Duchaine, and Adam Steel for helpful discussion about the project.

## Supplemental Information

**Supplemental Figure 1.**
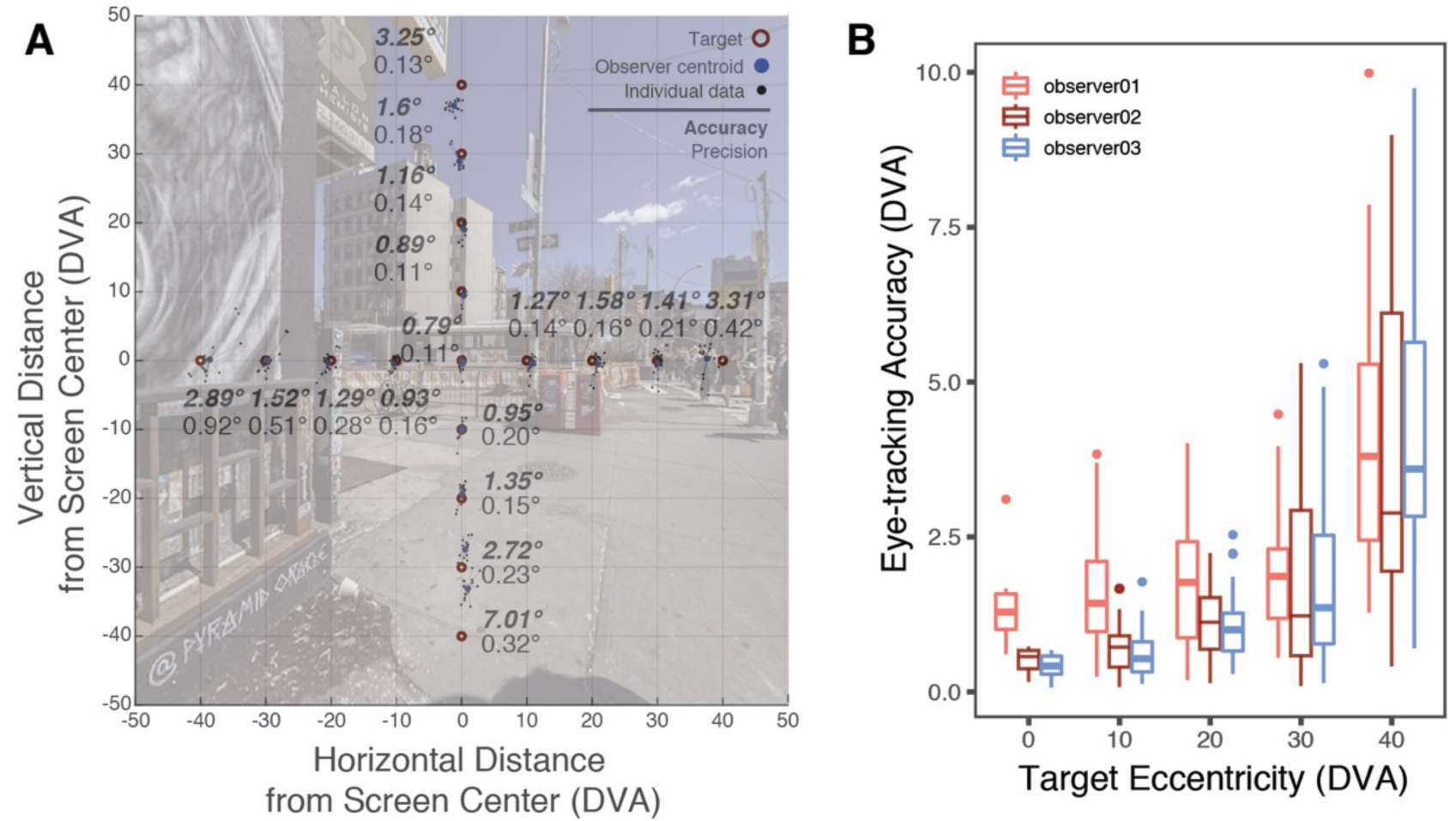
Gaze measured in 360 is accurate and precise. Three head-fixed participants made a series of saccades from a central fixation cross to 16 targets arranged in a cross and spanning the entire headmounted display’s field of view. Targets were located above, below, and to the right and left of screen center, and were located at distances of 10, 20, 30, and 40 DVA in each direction. Each observer completed this task six times, over separate sessions. The mean accuracy across observers and locations was 2.00 DVA +/− 0.38 STE, and the mean precision across observers and locations was 0.26 DVA +/− 0.05 STE. Eye-tracking accuracy decreased with eccentricity (F(1, 6.126) = 58.175, p < 0.001); however, gaze measured at screen center was accurate within <1 DVA, comparable to the reported accuracy of many mobile eye-tracking systems(Cognolato et al., 2018).

**Supplemental Figure 2.**
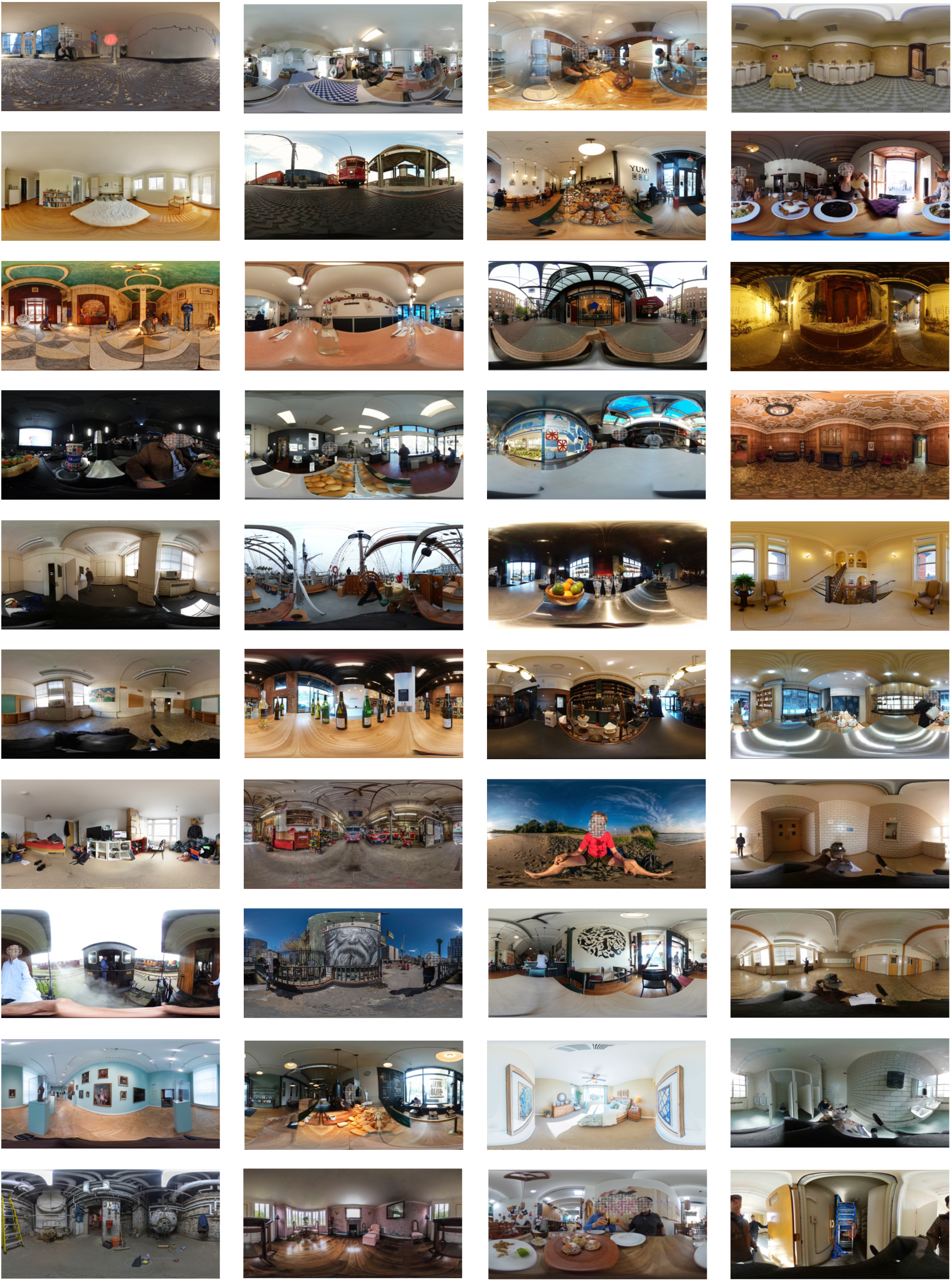
Experimental Stimuli.

**Supplemental Table 1:** Control analysis results summary. Meaning:salience:condition interaction remains significant when restricting analysis to the fields of view containing regions ranked in the top 50^th^ percentile for meaning.

|  | NumDF | DenDF | *F* value | *p* value |
| --- | --- | --- | --- | --- |
| condition | 1 | 5935512 | 6.78E+03 | < 0.001 |
| meaning | 1 | 5935478 | 6.89E+04 | < 0.001 |
| salience | 1 | 5935534 | 1.98E+04 | < 0.001 |
| condition:meaning | 1 | 5935512 | 2.21E+04 | < 0.001 |
| condition:salience | 1 | 5935512 | 3.39E+00 | 0.0656 |
| meaning:salience | 1 | 5935548 | 6.07E+04 | < 0.001 |
| condition:equator | 2 | 5935532 | 4.37E+05 | < 0.001 |
| condition:meaning:salience | 1 | 5935512 | 4.34E+02 | < 0.001 |

**Supplemental Table 2:** Control analysis results summary. Meaning:salience:condition interaction remains significant when accounting for groups of fixations in the active condition whose combined duration exceeded 5 seconds.

|  | **NumDF** | **DenDF** | **F value** | **p value** |
| --- | --- | --- | --- | --- |
| condition | 1 | 6163960 | 1.46E+04 | < 0.001 |
| meaning | 1 | 6163922 | 8.20E+04 | < 0.001 |
| salience | 1 | 6163986 | 1.24E+04 | < 0.001 |
| condition:meaning | 1 | 6163960 | 2.71E+04 | < 0.001 |
| condition:salience | 1 | 6163960 | 3.76E+02 | < 0.001 |
| meaning:salience | 1 | 6163998 | 4.75E+04 | < 0.001 |
| condition:equator | 2 | 6163980 | 4.71E+05 | < 0.001 |
| condition:meaning:salience | 1 | 6163960 | 1.49E+03 | < 0.001 |

